# The circadian clock contributes to the long-term water use efficiency of Arabidopsis

**DOI:** 10.1101/583526

**Authors:** Noriane M. L. Simon, Calum A. Graham, Nicholas E. Comben, Alistair M. Hetherington, Antony N. Dodd

## Abstract

In plants, water use efficiency is a complex trait derived from numerous physiological and developmental characteristics. Here, we investigated the involvement of circadian regulation in long-term water use efficiency. Circadian rhythms are generated by the circadian oscillator, which provides a cellular measure of the time of day. In plants, the circadian oscillator contributes to the regulation of many aspects of physiology, including stomatal opening, the rate of photosynthesis, carbohydrate metabolism and developmental processes. We investigated in Arabidopsis the impact of the misregulation of genes encoding a large number of components of the circadian oscillator upon whole plant, long-term water use efficiency. From this, we identified a role for the circadian oscillator in water use efficiency. This appears to be due to contributions of the circadian clock to the control of transpiration and biomass accumulation. We also identified that the circadian oscillator within guard cells can contribute to long-term water use efficiency. Our experiments indicate that knowledge of circadian regulation will be important for developing future crops that use water more efficiently.

**One-sentence summary:** The circadian clock in Arabidopsis makes an important contribution to long-term water use efficiency.

## Introduction

World population growth is increasing the demand for fresh water for agriculture, with climate change predicted to exacerbate this competition for water resources (Ruggiero et al., 2017). One strategy to sustainably increase agricultural production involves the improvement of crop water use (Condon et al., 2004; Xoconostle-Cazares et al., 2010; Hu and Xiong, 2014; Ruggiero et al., 2017). Because up to 97% of water taken up from the soil by plants is lost through stomatal transpiration (Yoo et al., 2009; Na and Metzger, 2014), the manipulation of transpiration represents an excellent candidate for designing crops with increased water use efficiency (Bertolino et al., 2019).

Plant water loss can be manipulated through changes in the regulation of stomatal opening and by altering stomatal density and patterning (Pei et al., 1998; Hugouvieux et al., 2001; Schroeder et al., 2001; Hetherington and Woodward, 2003; Yoo et al., 2010; Lawson and Blatt, 2014; Franks et al., 2015; Caine et al., 2019). In addition to stomatal responses to environmental cues such as light, temperature and phytohormones, there are circadian rhythms of stomatal opening (Gorton et al., 1989; Hennessey and Field, 1991). Circadian rhythms are self-sustaining biological cycles with a period of about 24 h. These rhythms are thought to adapt plants to daily cycles of light and dark, by anticipating daily changes in the environment and co-ordinating cellular processes. In higher plants, circadian rhythms are generated by several interlocked transcription-translation feedback loops known as the circadian oscillator (Hsu and Harmer, 2014). The phase of the circadian oscillator is adjusted continuously to match the phase of the environment through the process of entrainment, in response to light, temperature and metabolic cues (Somers et al., 1998; Millar, 2004; Salomé and McClung, 2005; Haydon et al., 2013; Webb et al., 2019). Additionally, the circadian oscillator communicates an estimate of the time of day to circadian-regulated features of the cell, initially through transcriptional regulation (Harmer et al., 2000). The known circadian oscillator controls circadian rhythms of stomatal opening because mutations that alter the circadian period or cause circadian arrhythmia lead to equivalent alterations in the circadian rhythm of stomatal opening (Somers et al., 1998; Dodd et al., 2004; Dodd et al., 2005). The circadian oscillator is also involved in the responses of guard cells to environmental cues such as drought and low temperature (Dodd et al., 2006; Legnaioli et al., 2009).

Circadian rhythms are often studied under conditions of constant light. However, the circadian oscillator is also important for the regulation of stomatal opening under cycles of light and dark. For example, constitutive overexpression of the circadian oscillator component CIRCADIAN CLOCK ASSOCIATED1 (CCA1; CCA1-ox) alters the daily regulation of stomatal opening such that stomatal conductance increases steadily throughout the photoperiod (Dodd et al., 2005). In comparison, stomatal conductance in wild type plants remains relatively uniform during the photoperiod and is lower than in CCA1-ox (Dodd et al., 2005). Similarly, guard cell-specific overexpression of CCA1 generally causes greater stomatal opening during the light period, and alters drought response phenotypes (Hassidim et al., 2017). Modelling suggests that under light/dark cycles, the circadian oscillator contributes at the canopy scale to daily rhythms in stomatal aperture and carbon assimilation in bean and cotton (Resco de Dios et al., 2016).

The contribution of the circadian oscillator to both stomatal opening and biomass accumulation (Dodd et al., 2005; Graf et al., 2010) suggests that the circadian oscillator might make an important contribution to long-term water use efficiency (WUE). WUE is the ratio of carbon dioxide incorporated through photosynthesis into biomass to the amount of water lost through transpiration. At the single leaf level, instantaneous, intrinsic WUE is often measured with gas exchange techniques and expressed as net CO_2_ assimilation per unit of water transpired (Vialet-Chabrand et al., 2016; Ruggiero et al., 2017; Ferguson et al., 2018). However, such measurements do not provide an accurate representation of WUE over the plant lifetime, which is influenced by features such as leaf position, dark respiration, and time of day changes in instantaneous WUE (Condon et al., 2004; Tomás et al., 2014; Medrano et al., 2015; Ferguson et al., 2018). It is important to note that high WUE under well-watered conditions is not the same as drought resistance, because drought resistance relates to the capacity to maintain transpirational water supply under water-limited conditions through strategies such as expanded root systems (Blum, 2009). This means that WUE does not correlate reliably with drought resistance (Kobata et al., 1996).

Given that the circadian oscillator affects stomatal opening and biomass accumulation (Gorton et al., 1989; Hennessey and Field, 1991; Dodd et al., 2005; Edwards and Weinig, 2010; Graf et al., 2010; Edwards et al., 2012), we hypothesized that specific components of the circadian oscillator might make an important contribution to the long-term WUE of plants. Although the circadian oscillator influences stomatal opening and biomass accumulation, the influence of the circadian oscillator upon long-term WUE of plants remains unknown. This is an important question for understanding roles for circadian regulation in crops, because the long-term water use efficiency ultimately determines the amount of water that is required for a given yield of the crop. Therefore, we investigated the impact of the misregulation of parts of the circadian oscillator upon the long-term WUE of Arabidopsis. We identified that the circadian oscillator has profound effects upon the long-term WUE of plants. Importantly, some alterations in oscillator function increase long-term WUE, suggesting potential targets for future improvements of crop WUE.

## Results

### Circadian oscillator components contribute to long-term water use efficiency

We identified that correct regulation of genes encoding circadian oscillator components makes an important contribution to WUE. 32 single mutants or overexpressors of genes associated with circadian regulation, representing 21 circadian oscillator-associated components, were surveyed for WUE alterations (Fig. 1) using a previously-described method (Wituszyńska et al., 2013). 44 % of the mutants or overexpressors examined had a significantly different WUE from the wild type (Fig. 1, Table 1). The *elf3*-1, *prr9*-1, *tps1*-11, *tps1*-12, and *ztl*-1 mutants, as well as CCA1, TOC1 and KIN10 overexpressors, had significantly lower WUE than the wild type (Fig. 1, Table 1; Table S1). The *gi*-2, *gi*-11, *grp7*-1, *tej*-1 and *tic*-2 mutants had significantly greater WUE than the wild type (Fig. 1, Table 1; Table S1). This suggests that misregulating the expression of circadian clock components CCA1, ELF3, GI, GRP7, PRR9, TEJ, TIC, TOC1 and ZTL can change whole plant long-term WUE (Fig. 1, Table 1). WUE was also altered by changing the expression of the energy signalling pathway components TPS1 and KIN10 that also provide inputs to the circadian oscillator (Shin et al., 2017; Frank et al., 2018) (Fig. 1, Table 1). Overall, these data show that correct expression of a variety of circadian clock-associated genes contributes to long-term WUE of Arabidopsis.

**Figure 1.**
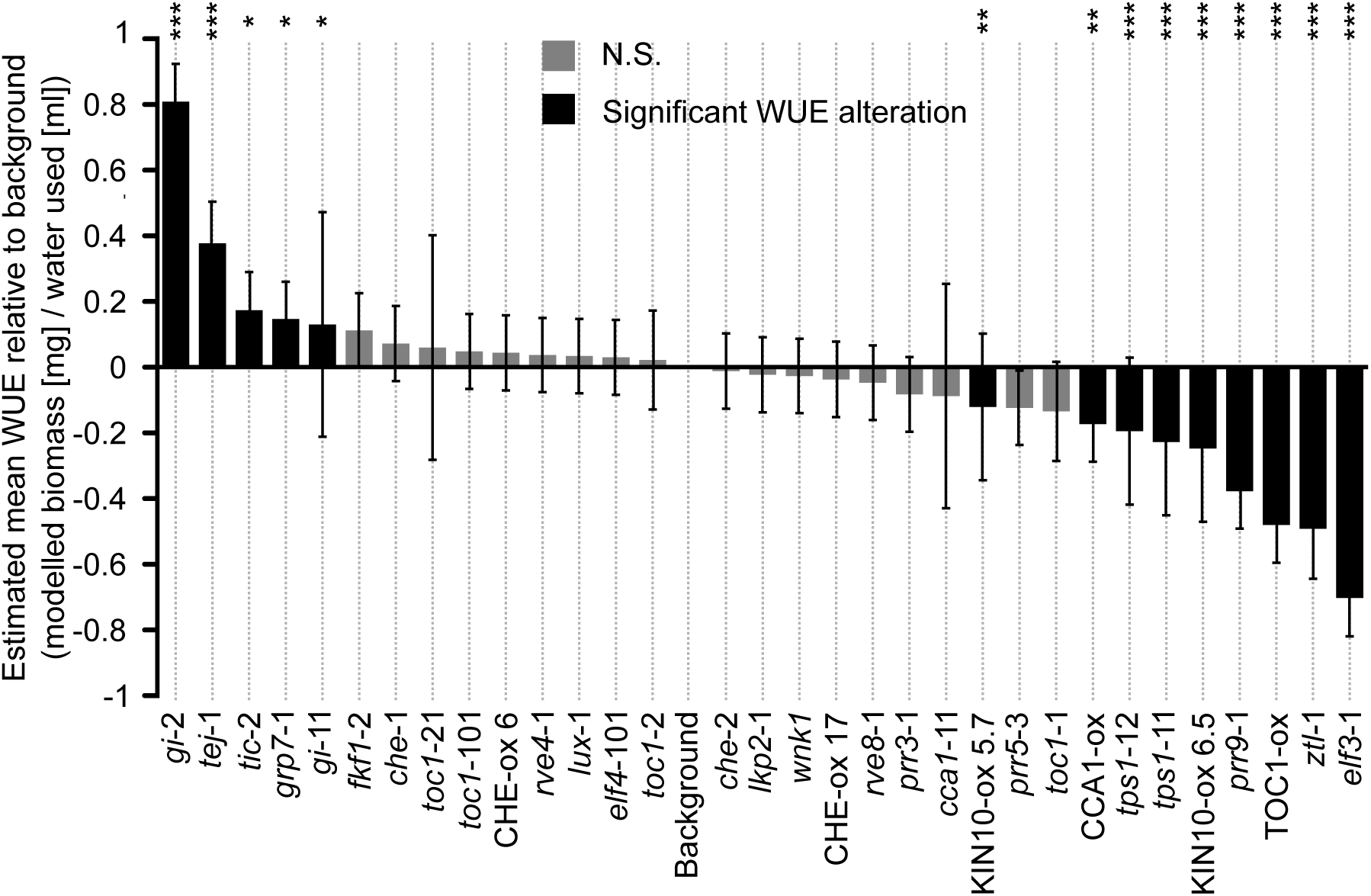
The circadian clock regulates long-term water use efficiency of Arabidopsis under light/dark cycles. Estimated WUE of each genotype relative to the estimated WUE of its background, derived from linear mixed effects models that combine all 18 separate batches of experimentation, each having 5 - 15 replicate plants per genotype. Statistical analysis derived from pairwise post-hoc comparisons between mutants and corresponding backgrounds, using the Kenward-Roger method for determining degrees of freedom and Tukey method for P-value adjustment. s.e.m. reflects modelled data, rather than the range of values sampled from the plants.

**Table 1.**
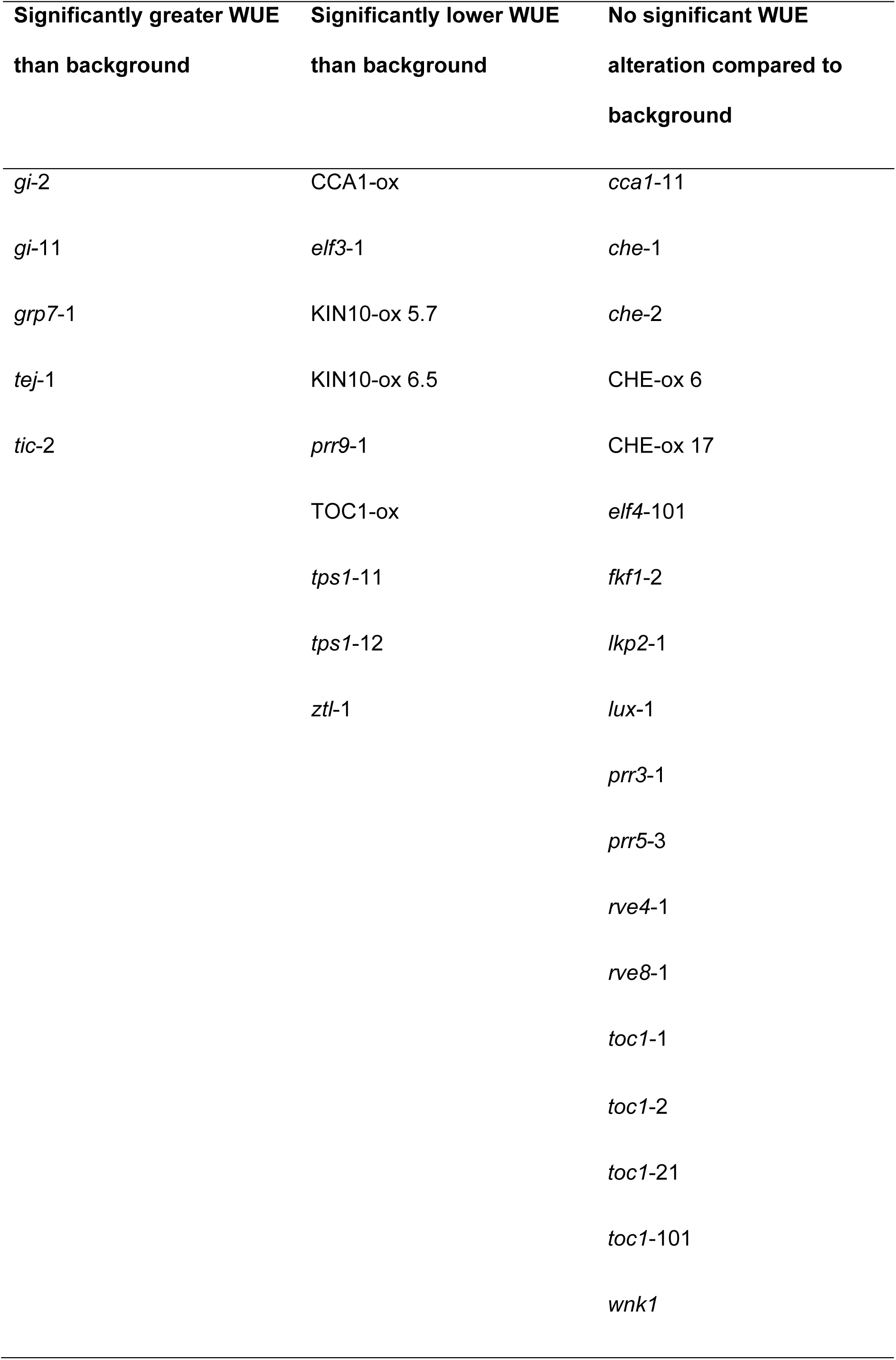
Genotypes with misregulated circadian clock-associated genes that have altered long-term WUE. Findings are summarized from Fig. 1 and derived from statistical significance within linear mixed effects models.

Within this experiment, each background accession had a distinct WUE (C24: 3.01 ± 0.07 mg ml^−1^; Col-0: 2.22 ± 0.02 mg ml^−1^; L. *er.*: 1.60 ± 0.04 mg ml^−1^; Ws: 1.91 ± 0.06 mg ml^−1^) (Fig. S2). These differences between WUE of different Arabidopsis accessions are consistent with previous studies of WUE, stomatal function and stomatal density in Arabidopsis (Nienhuis et al., 1994; Woodward et al., 2002; Dodd et al., 2004; Masle et al., 2005; Karaba et al., 2007; Ruggiero et al., 2017; Ferguson et al., 2018).

We hypothesised that variations in WUE might be associated with specific circadian phenotypes in the mutants and overexpressors that we tested. For example, mutations in circadian clock genes expressed with a particular phase (e.g. morning-expressed or evening-expressed genes) might have a more pronounced effect on WUE. Likewise, the nature of the circadian period change or flowering time change resulting from misexpression of each oscillator component might be associated with certain changes in WUE. To test this, we compared the data from our WUE screen with the circadian phase of expression of each mutated or overexpressed gene. We also compared the direction of change of WUE to the period and flowering time phenotypes that arise from each mutant or overexpressor (Hicks et al., 1996; Fowler et al., 1999; Schultz et al., 2001; Doyle et al., 2002; Nakamichi et al., 2002; Yanovsky and Kay, 2002; Imaizumi et al., 2003; Más et al., 2003; Murakami et al., 2004; Farré et al., 2005; Hazen et al., 2005; Baena-González et al., 2007; Ding et al., 2007; Niwa et al., 2007; Streitner et al., 2008; Wang et al., 2008; Baudry et al., 2010; Nakamichi et al., 2010; Rawat et al., 2011; Wahl et al., 2013; Hsu and Harmer, 2014). We note that the phenotypes reported by these studies were often identified under constant conditions, with flowering time experiments performed under short or long photoperiods, whereas our experiments occurred under cycles of 8 h light / 16 h darkness.

There was no obvious relationship between the circadian phenotypes reported to arise from each mutant or overexpressor investigated and the WUE of each of these lines (Fig. 2A, B, C). For example, mutating night-phased oscillator components can either decrease or increase WUE (Fig. 2A). Mutants that cause long circadian periods and short circadian periods can both increase and decrease WUE, although WUE was unaltered in the mutants that are reported to not alter the period (Fig. 2B). Furthermore, mutants and overexpressors that cause both early and delayed flowering can each increase and decrease WUE (Fig. 2C).

**Figure 2.**
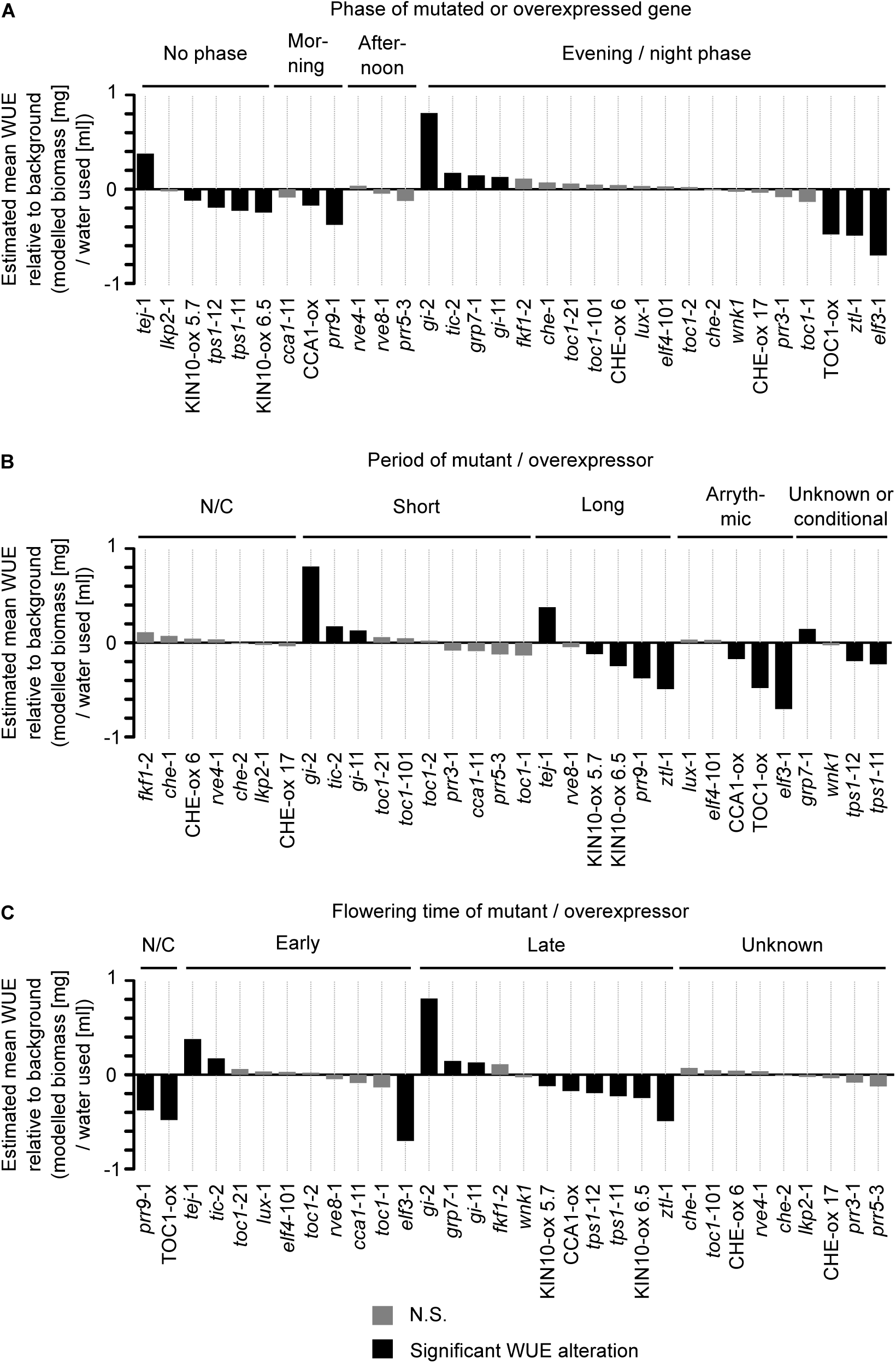
Relationship between circadian clock-associated characteristics and water use efficiency. (A-C) Estimated WUE grouped according to (A) phase of expression of each mutated or overexpressed gene, and the (B) period or (C) flowering time alteration caused by mutation or overexpression of each gene indicated. Shading of bars on graphs indicates statistical significance. Studies describing the phase of expression, period and flowering time of the genotypes tested are identified in the main text. N/C indicates no change. We note that the phase of expression and period data used for this analysis were often identified in previous studies under constant conditions, in contrast to our experiments under light/dark cycles. Estimated WUE relative to its corresponding background is derived from linear mixed effects models combining all 18 separate batches of experimentation, each having 5 - 15 replicate plants per genotype. Statistical analysis derived from pairwise post-hoc comparisons between mutants and corresponding backgrounds, using the Kenward-Roger method for determining degrees of freedom and Tukey method for P-value adjustment.

We were interested to determine whether the WUE alterations caused by misregulation of circadian oscillator gene expression arose from changes in either biomass accumulation or transpiration. Genotypes with low biomass accumulation generally had lower water use (Fig. 3A, B). The results for *tej*-1 (Fig. 3A, B) should be treated with caution because the model fits were very poor (Table S1). No mutants or overexpressors tested increased the biomass relative to the corresponding background genotype (Fig 3A). Together, this indicates that the altered WUE phenotypes in some genotypes with misregulated circadian clocks was not due to an alteration in just one of either water use or biomass accumulation (Fig. 3A, B). Instead, the altered WUE of lines with misregulated circadian clock genes appears to be due to the net effect of altered biomass accumulation and altered transpiration in these genotypes (Fig. 3A, B).

**Figure 3.**
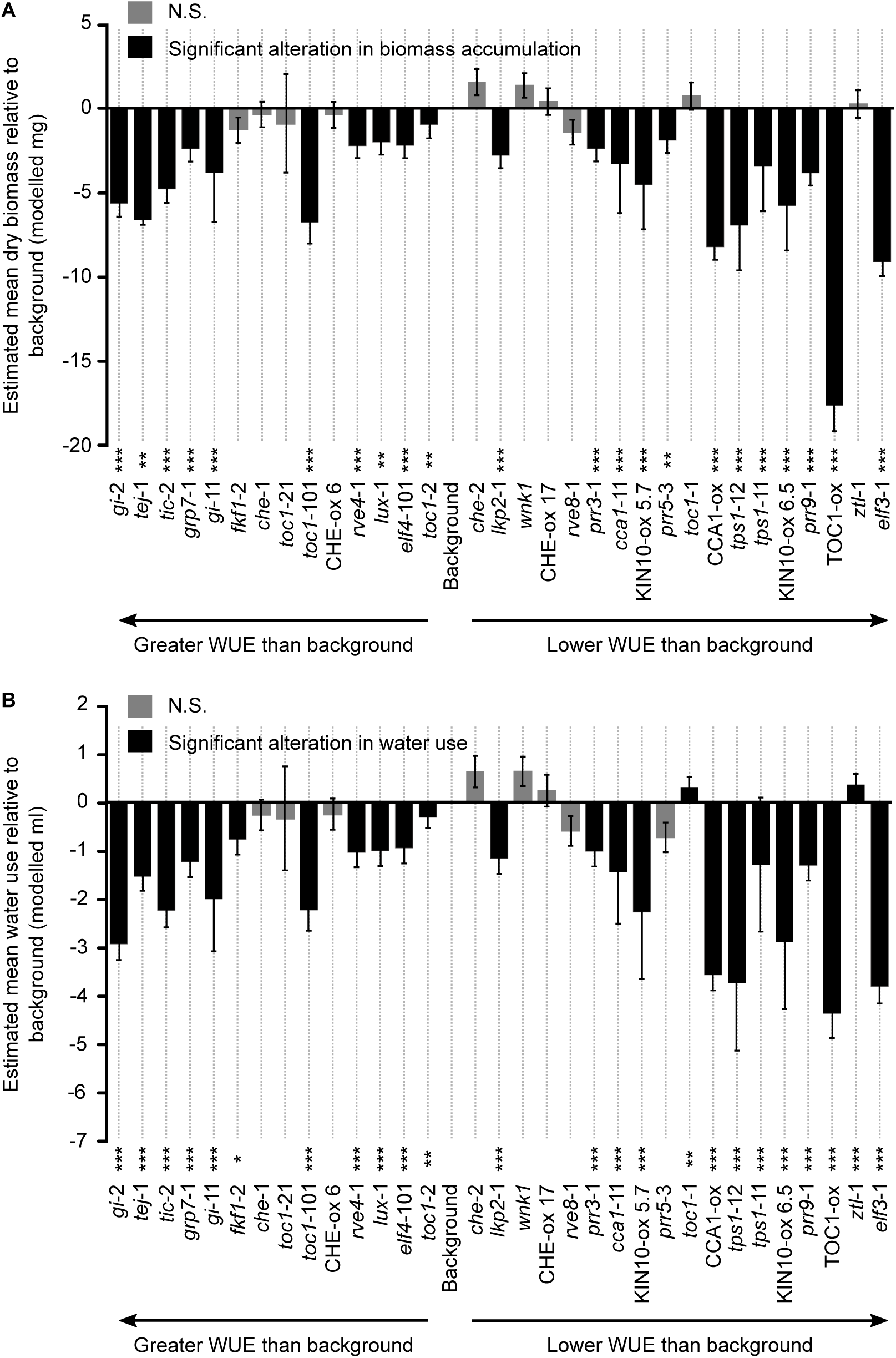
Manipulating the expression of genes associated with circadian regulation alters estimated WUE by changing both water use and biomass accumulation. (A) Biomass accumulation loss and (B) water loss for each genotype relative to its respective background over the course of the experiments. Genotypes are ordered according to their WUE calculated in Fig. 1. Data are derived from linear mixed effects models that combine all 18 separate batches of experimentation, each having 5 - 15 replicate plants per genotype. Statistical analysis derived from pairwise post-hoc comparisons between mutants and corresponding backgrounds, using the Kenward-Roger method for determining degrees of freedom and Tukey method for P-value adjustment. Note the model fit for the *tej*-1 mutant was poor (Table S1). Genotypes are ordered from greatest (left) to lowest (right) estimated WUE. s.e.m. reflects modelled data, rather than the range of values sampled from the plants.

### Circadian regulation of water use efficiency combines multiple traits

Mutation or overexpression of components of the circadian oscillator can cause changes in the development of Arabidopsis, such as alterations in rosette size, leaf shape and petiole length (Fig. 4A) (Zagotta et al., 1992; Schaffer et al., 1998; Wang and Tobin, 1998; Dodd et al., 2005; Ruts et al., 2012; Rubin et al., 2018). These changes are likely to have implications for gas exchange because, for example, spatially separated leaves are predicted to transpire more water (Bridge et al., 2013). We investigated whether the changes in WUE that were identified by our screen might arise from differences in rosette architecture between the circadian clock-associated mutants and overexpressors and the corresponding backgrounds. There was a weak positive correlation between rosette leaf surface area and WUE (*r* = 0.400; *r^2^* = 0.160; *p* < 0.001) (Fig. 4B). Therefore, this suggests that approximately 16% of variability in WUE can be explained by the variations in rosette leaf surface area that arise from misregulation of the circadian oscillator.

**Figure 4.**
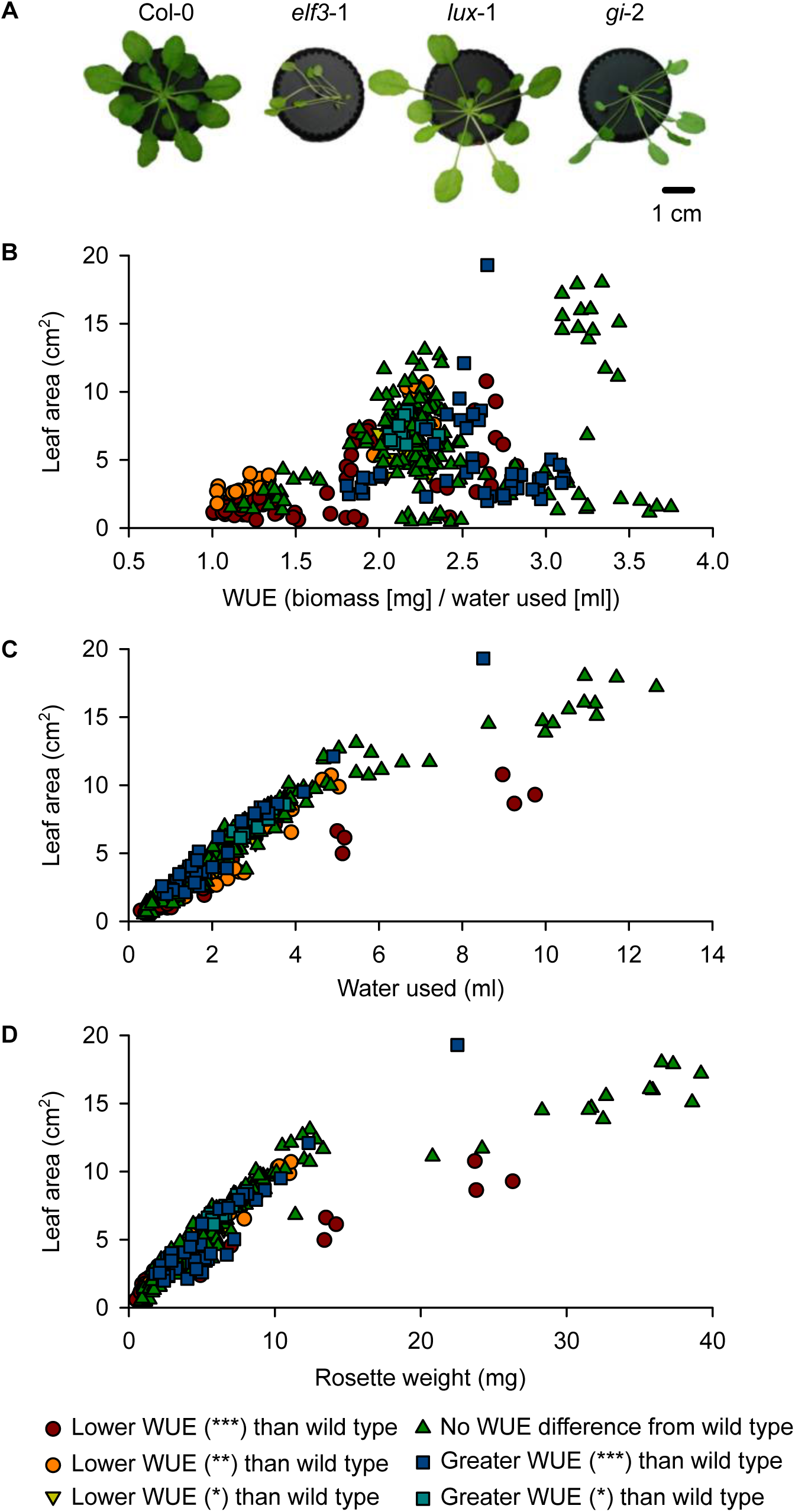
The circadian oscillator alters WUE partially by changing rosette architecture. (A) Altering circadian-associated gene expression can affect rosette architecture and size, as illustrated for *elf3*-1, *lux*-1, and *gi*-2 in the Col-0 background. Image backgrounds removed for clarity. Variation in rosette leaf surface area across the genotypes investigated explained (B) 16% of variation in WUE (*p* < 0.001, *r* = 0.400, *r*^2^ = 0.160), (C) 83% of variation in transpiration (*p* < 0.001, *r* = 0.912, *r*^2^ = 0.832) and (D) 73% of variation in rosette dry biomass (*p* < 0.001, *r* = 0.857, *r^2^* = 0.734). Data were analysed using Pearson correlation tests.

In comparison, rosette leaf surface area was strongly correlated with each of the individual parameters of water used and dry biomass accumulated. The variation in rosette surface area accounted for 83% of the variability in water transpired across the genotypes (Fig. 4C). Furthermore, the variation in rosette surface area accounted for 73% of the variability in biomass accumulation across the genotypes (Fig. 4D), which is unsurprising given that larger leaves are likely to contain more biomass.

This demonstrates that one way that circadian regulation affects WUE is through the influence of the circadian oscillator upon plant development and rosette architecture, but this variation in leaf area does not account for the majority of the influence of circadian regulation upon WUE. It also further supports the notion that the influence of the circadian oscillator upon WUE is complex, and cannot be explained by variation in one of water use or biomass accumulation alone.

### Contribution of circadian regulation in guard cells to water use efficiency

Next, we investigated whether the circadian oscillator within guard cells contributes to long-term WUE. There is evidence that guard cells contain a circadian oscillator that regulates stomatal opening (Gorton et al., 1989; Hassidim et al., 2017). To investigate the contribution of the guard cell circadian oscillator to WUE, we overexpressed two circadian oscillator components (*CCA1*, *TOC1*) in guard cells, using two guard cell-specific promoters (*GC1*, *MYB60*) for each of *CCA1* and *TOC1* (Fig. 5A) (Cominelli et al., 2005; Galbiati et al., 2008; Yang et al., 2008; Nagy et al., 2009; Meyer et al., 2010; Cominelli et al., 2011; Bauer et al., 2013; Rusconi et al., 2013). *GC1* is a guard cell-specific promoter that is relatively unresponsive to a variety of environmental cues (cold, light, ABA, gibberellin) (Yang et al. 2008). We used the full-length *MYB60* promoter sequence, because truncated and chimeric versions of this promoter appear to have weaker activity and/or become rapidly downregulated by dehydration and ABA (Francia et al., 2008; Cominelli et al., 2011; Rusconi et al., 2013). This produced four sets of transgenic lines; *GC1::CCA1:nos* (*GC*), *GC1::TOC1:nos* (*GT*), *MYB60::CCA1:nos* (*MC*) and *MYB60::TOC1:nos* (*MT*) (Fig. 5A). We termed these guard cell specific (GCS) plants. We confirmed the guard cell specificity of the *GC1* and *MYB60* promoters in our hands, by driving green fluorescent protein (GFP) under the control of these promoters. GFP accumulation was restricted to the guard cells (Fig. S3A, B). There was not a circadian oscillation in the activity of either the *GC1* or *MYB60* promoter under our experimental conditions (Fig. S3C), demonstrating that these promoters were appropriate for constitutive overexpression of circadian oscillator components within guard cells in our experiments.

**Figure 5.**
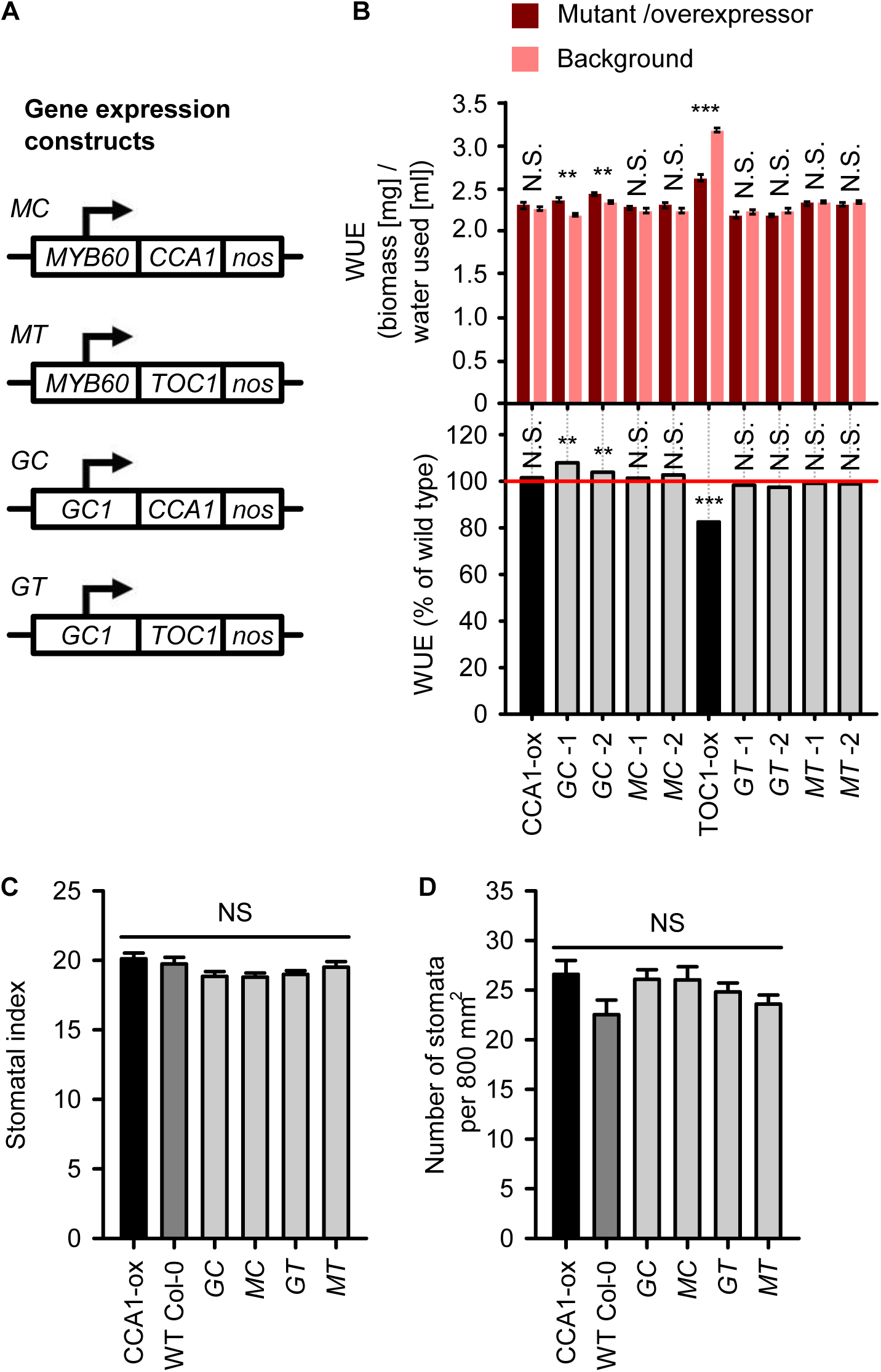
Long-term WUE can be altered by misexpression of circadian oscillator genes within stomatal guard cells. (A) Constructs used to overexpress *CCA1* or *TOC1* coding sequence under control of *GC1* or *MYB60* promoters. (B) Guard cell *CCA1* overexpression can increase WUE. WUE expressed as absolute WUE and percentage of the wild type (normalised to wild type as 100%, red reference line). Data were analysed by one-way ANOVA (p < 0.001) followed by pairwise post-hoc Tukey comparisons between transgenic lines and their corresponding backgrounds (** = *p* < 0.01; *** = *p* < 0.001; N.S. = *p* > 0.05; *n* = 5 - 15). (C, D) Guard cell *CCA1* or *TOC1* overexpression does not affect (C) stomatal index nor (D) stomatal density. Data were analysed with ANOVA and Tukey’s post hoc tests (N.S. = *p* > 0.05; *n* = 19 - 32; mean ± S.E.M.). Bar colours identify the whole plant overexpressor control (black), wild type control (dark grey), and guard cell-specific overexpressor genotypes (light grey).

To further verify the guard cell-specific overexpression of *CCA1* and *TOC1* in the GCS plants, we examined *CCA1* and *TOC1* transcript accumulation within guard cells. Under constant light conditions, we measured *CCA1* transcript accumulation in epidermal peels at dusk (when *CCA1* transcript abundance is normally low in the wild type) and *TOC1* transcript accumulation at dawn (when *TOC1* transcript abundance is normally low in the wild type). Guard cell *CCA1* overexpressors had greater *CCA1* transcript abundance in epidermal peels at dusk than the wild type (Fig. S3D), and guard cell *TOC1* overexpressors had greater *TOC1* transcript abundance at dawn than the wild type (Fig. S3D). These data indicate that *CCA1* and *TOC1* were overexpressed within the guard cells of the guard cell-specific *CCA1* or *TOC1* overexpressor plants that we generated, respectively.

We investigated the effect on WUE of overexpression of CCA1 and TOC1 within guard cells. Two independent *GC1::CCA1* lines (*GC*-1 and *GC*-2) were significantly more water use efficient than the wild type (*GC*-1: *p* < 0.001; *GC*-2: *p* = 0.002) (Fig. 5B). *GC*-1 and *GC*-2 were 8% and 4% more water use efficient than the wild type, respectively (Fig. 5B). In comparison, two independent *MYB60::CCA1* did not have greater WUE than the wild type (*p* > 0.05) (Fig. 5B). This suggests that overexpressing *CCA1* in guard cells can increase whole plant long-term WUE in a promoter-specific manner. Overexpression of *TOC1* in guard cells with both the *GC1* and *MYB60* promoters did not alter WUE (*p* > 0.05) (Fig. 5B).

This suggests that decreased WUE in constitutive TOC1-ox plants (Fig. 1, Fig. 5B) might not be explained by overexpression of TOC1 within the guard cells, and that this decreased WUE might instead be due to TOC1 overexpression in other cell types. In this particular experiment we did not see an alteration in CCA1-ox WUE relative to its background, contrasting Fig. 1. This could suggest that the WUE alteration in CCA1-ox is fairly small or variable. Because the stomatal density was unaltered relative to the wild type in the guard cell overexpressors of *CCA1* and *TOC1* (Fig. 5C, D), the WUE phenotypes that we identified from these lines might be caused by alterations in processes within guard cells, such as those regulating stomatal aperture, rather than altered stomatal density.

## Discussion

### Pervasive influence of the circadian oscillator upon water use efficiency

Our data indicate that the circadian oscillator is important for regulating the long-term WUE of Arabidopsis. Misregulation of several functional subsections of the circadian oscillator altered the WUE. Misexpression of morning (PRR9, CCA1), late day (GI) and evening (TOC1, ZTL, ELF3) components of the circadian oscillator all perturb WUE under our experimental conditions (Fig. 1). Additionally, mutation of TEJ and GRP7 alters WUE (Fig. 1). Therefore, oscillator components that impact WUE are not confined to a specific expression phase or architectural feature (e.g. morning loop) within the multi-loop circadian oscillator. Misexpression of genes encoding some proteins that provide environmental inputs to the circadian oscillator (ELF3, TPS1, ZTL, KIN10; (Covington et al., 2001; Kim et al., 2007; Shin et al., 2017; Frank et al., 2018)) also alters WUE (Fig. 1). Together, this suggests that the entire circadian oscillator can influence WUE, and that alterations in water use that are caused by mutations to the circadian oscillator are not confined to a specific sub-loop of the circadian oscillator, or restricted to its input or output pathways. One explanation for these circadian-system wide alterations in WUE relates to the nature of feedback within the circadian oscillator. The complex feedback and interconnectivity of the circadian oscillator means that individual components of the circadian oscillator that directly influence stomatal function or water use are likely to be altered by mutations that are distal to that component. Therefore, if correct circadian timing is required for optimum water use efficiency, multiple components of the circadian oscillator are likely to influence water use efficiency.

Alternatively, because mutation of a number of components of the circadian oscillator had no effect upon WUE, it is possible that the oscillator components that influence WUE do so through roles in directly regulating outputs of the circadian oscillator, such as by regulating genes involved in stomatal function.

The sugar signalling proteins TPS1 and KIN10 influence a broad range of phenotypes, in addition to participating in circadian entrainment (Baena-González et al., 2007; Gómez et al., 2010; Paul et al., 2010; Delatte et al., 2011; Shin et al., 2017; Frank et al., 2018; Nietzsche et al., 2018; Simon et al., 2018). The *tps1*-12 TILLING mutant of TPS1 decreases stomatal aperture and increases the ABA sensitivity of guard cells (Gómez et al., 2010), whereas we found that *tps1*-11 and *tps1*-12 had lower long-term WUE than the wild type (Fig. 1). Lower biomass accumulation in *tps1*-11 and *tps1*-12 (Fig. 3A) was consistent with slow growth of these alleles (Gómez et al., 2010). Overall, this suggests that the decreased stomatal aperture of *tps1*-12 mutants (Gómez et al., 2010) does not translate into an overall increase in WUE, perhaps due to slower growth or misregulated ABA signalling in the *tps* mutants (Gómez et al., 2010). The broad range of phenotypes that are altered in *tps1*-11, *tps1*-12 and KIN10-ox indicates that these genotypes might alter WUE through mechanisms other than circadian regulation.

### Potential roles for the evening complex in WUE

Our finding that ELF3 can influence WUE (Fig. 1) is supported by previous evidence. Under constant light conditions, wild type Arabidopsis has circadian rhythms of stomatal aperture, whereas *elf3* stomata are constantly open and unresponsive to light and dark (Kinoshita et al., 2011). Furthermore, ELF3 negatively regulates blue light-mediated stomatal opening (Kinoshita and Hayashi, 2011). Therefore, perturbation of the anticipation of day/night transitions or responses to environmental cues in *elf3* stomata might cause long-term alterations in WUE.

ELF3 binds to the *PRR9* promoter and *elf3*-1 has elevated PRR9 transcript abundance (Thines and Harmon, 2010; Dixon et al., 2011; Herrero et al., 2012). The low WUE of *elf3*-1 might potentially be caused by altered PRR9 expression, because misregulation of *PRR9* also affected WUE (Fig. 1). In a similar fashion, ELF3/ELF4 signalling represses PRR7, and *elf3*-1 has elevated *PRR7* transcript abundance (Herrero et al., 2012). Under light-dark cycles, *elf3*-1 also has high and constitutive *GI* expression (Fowler et al., 1999), and *elf3*-1 and *gi* mutants have opposite WUE phenotypes (Fig. 1). Therefore, the WUE phenotype of *elf3*-1 (Fig. 1) might be caused by disruption of ELF3 itself, or specific perturbations of PRR7, PRR9 and/or GI expression.

Mutating further components of the evening complex (EC) (ELF4 and LUX) did not affect WUE (Fig. 1). This is despite these genes influencing circadian oscillator function and plant physiology (Hsu and Harmer, 2014; Huang and Nusinow, 2016), and nocturnal regulation of stomatal aperture altering WUE (Costa et al., 2015; Coupel-Ledru et al., 2016). One possibility is that the impact of *elf3-*1 on WUE may be greater than that of *elf4* or *lux* because ELF3 is key to EC scaffolding, with ELF3 operating genetically downstream from ELF4 and LUX (Herrero et al., 2012; Huang and Nusinow, 2016).

The EC binds upstream of and regulates a variety of other genes that might also underlie the WUE alterations in *elf3-*1 mutants (Ezer et al., 2017). This includes regulators of growth, components of the photosynthetic apparatus, and genes associated with phytohormone signalling. This means that potential roles for the EC in WUE might occur through several physiological mechanisms. There also appears to be a negative relationship between temperature and EC promoter binding (Ezer et al., 2017), so it is possible that any influence of the EC upon WUE might be temperature-sensitive.

ELF4 appears to play a greater role in circadian regulation in the vascular tissue than stomatal guard cells, with vasculature expression up to ten times higher than other tissues (Endo et al., 2014). Processes within the vasculature can affect WUE; for example, mutations in *CELLULOSE SYNTHASE CATALYTIC SUBUNIT7* (*CESA7*) might impact water use through effects of the collapse of the vasculature upon guard cell size (Liang et al., 2010). Because *elf3*-1 affects WUE differently from *elf4*-101 and *lux*-1 (Fig. 1), ELF3 might regulate WUE independently from ELF4 and LUX.

### Multiple physiological causes of altered WUE in circadian oscillator mutants

Our data suggest that changes in WUE caused by misexpression of circadian clock components might be due to a combination of physiological factors. Many mutants or overexpressors tested alter both biomass accumulation and water loss, often in the same direction (Fig. 3A, B), so mutations to the circadian oscillator did not alter water use by specifically altering either carbon assimilation or transpiration. This is consistent with previous work demonstrating that both stomatal opening and CO_2_ fixation is perturbed in circadian arrhythmic plants under light/dark cycles (Dodd et al., 2005), and also with the finding that daily carbohydrate management is dependent upon correct circadian regulation (Graf et al., 2010). We speculate that delayed or advanced stomatal and photosynthetic responses to the day-night cycle might occur in circadian period mutants, because period mutants inaccurately anticipate the onset of dawn (Dodd et al., 2014). Circadian clock mutants might also affect WUE by changing the sensitivity of stomatal movements and photosynthesis to environmental transitions, because there is circadian gating of the responses of both stomata and photosynthesis to environmental cues (Dodd et al., 2006; Kinoshita et al., 2011; Litthauer et al., 2015; Joo et al., 2017; Cano-Ramirez et al., 2018). Some effects of the circadian oscillator upon WUE arise from alterations in leaf size that occur in some circadian oscillator mutants (Fig. 4A, B). This suggests that developmental alterations arising from lesions in the circadian oscillator can lead to changes in WUE. Such developmental alterations might alter WUE by changing airflow around the rosette, boundary layer conductance, or internal leaf structure.

It has been reported previously that during the light period of light/dark cycles, CCA1-ox has greater stomatal conductance than the wild type and decreased CO_2_ assimilation and biomass accumulation (Dodd et al., 2005; Graf et al., 2010). If these alterations in growth, CO_2_ fixation and transpiration persist throughout the vegetative growth phase, it might be predicted that CCA1-ox would have lower long-term WUE than the wild type. We found that both biomass accumulation and water loss were reduced significantly in CCA1-ox relative to its background (Fig. 3A, B), with the ratio between the two parameters indicating also a significant decrease in WUE of CCA1-ox (Fig. 1). This might be due to alterations in gas exchange reported previously (Dodd et al., 2005), and also other developmental changes caused by CCA1 overexpression.

### Contribution of circadian regulation in guard cells to water use efficiency

We also investigated whether the circadian oscillator within guard cells contributes to long-term WUE. This involved overexpressing two circadian clock genes in guard cells using two different guard cell-specific promoters. Comparable approaches have been adopted to investigate roles of specific cell types in the functioning of the circadian system and their relationships with physiology and development (Endo et al., 2014; Shimizu et al., 2015; Hassidim et al., 2017). Under our experimental conditions, we did not identify consistent alterations in the long-term WUE of seedlings overexpressing *CCA1* or *TOC1* in stomatal guard cells (Fig. 5B). This could indicate that decreased long-term WUE of TOC1-ox plants (Fig. 1) arises from altered circadian regulation in cell types other than guard cells. Whilst two lines harbouring a *GC1::CCA1* construct had greater WUE than the wild type, WUE was unaltered in comparable lines harbouring *MYB60::CCA1* (Fig. 5B). The differing WUE phenotype of *GC1::CCA1* and *MYB60::CCA1* might be explained by differences in promoter strength, because the *GC1* promoter appears to have somewhat greater activity than the *MYB60* promoter (Fig. S3D, E). Although both promoters are guard cell-specific in our hands (Fig. S3), we cannot exclude the possibility of ectopic promoter activity.

Interestingly, *GC1::CCA1* is reported to have greater drought sensitivity of long-term biomass accumulation than the wild type (Hassidim et al., 2017), whereas we found that *GC1::CCA1* had greater WUE than the wild type (Fig. 5B). This might reflect the integration of circadian regulation into ABA signalling (Legnaioli et al., 2009; Robertson et al., 2009), or occur because guard cell circadian regulation is required for correct guard cell metabolism and/or stomatal movements under conditions of abiotic stress. For example, circadian regulation is proposed to participate in daily cycles of triacylglycerol mobilization that are important for stomatal opening (McLachlan et al., 2016). Together, these findings suggest that guard cell circadian regulation is important under both well-watered conditions and conditions of environmental stress (Fig. 5C) (Robertson et al., 2009; Hassidim et al., 2017), with circadian regulation in other tissues also contributing to overall WUE. It would be informative in future to perform reverse genetic screening of the dehydration tolerance or long-term drought tolerance of sets of circadian clock mutants. However, because well-watered WUE is not a drought tolerance trait (Blum, 2009), it possible that different circadian clock mutant alleles might confer dehydration or drought tolerance compared with those alleles that alter WUE (Fig. 1).

### Conclusions

We show that circadian regulation contributes to whole plant long-term WUE under cycles of day and night. This control occurs partly through the influence of components of the circadian oscillator upon rosette architecture. Mutation or overexpression of CCA1, TOC1, ELF3, GI, GRP7, PRR9, TEJ, TIC and ZTL altered WUE under our experimental conditions. The roles of these genes in WUE may be independent or overlapping, and their WUE phenotypes might be due to direct effects of these genes, or indirect effects on transcript and/or protein abundance of other circadian clock gene(s). Misregulation of the expression of CHE, FKF1, LKP2, RVE4, RVE8, PRR3, PRR5, ELF4, LUX and WNK1 did not appear to alter WUE under our experimental conditions.

Our results have broad implications. Firstly, our data suggest that alterations in circadian function that arise during crop breeding could have the potential to alter WUE. Therefore, manipulation of the functioning of the circadian oscillator might represent a pathway to tune the WUE of crops. Second, our results indicate that circadian regulation in a single cell type can have implications for whole-plant physiology. Third, our experiments with guard-cell misregulation of circadian oscillator genes suggest that the circadian oscillator within guard cells and other cell types influences WUE. Finally, our findings suggest that circadian regulation potentially alters a single trait (WUE) by affecting many aspects of physiology, along with leaf area. Overall, our study demonstrates that the circadian oscillator is important for the water use efficiency of Arabidopsis plants during their entire vegetative growth period. In future, it will be informative to distinguish the contribution to overall WUE of circadian regulation within additional cell types, such as the mesophyll, vascular tissue, and root cell types. It will also be important to identify specific mechanisms underlying the WUE phenotypes, and determine the extent to which these findings scale to crop species.

## Materials and methods

### Plant material and growth conditions

Arabidopsis (*Arabidopsis thaliana* (L.) Heynh.) seeds were surface-sterilised as described previously (Noordally et al., 2013). For experiments investigating stomatal density and index, seeds were stratified for 3 days at 4 °C, then sown on compost mix comprising a 3:1 ratio of coarsely sieved Levington Advance F2 seed compost (Everris) and horticultural silver sand (Melcourt), supplemented with 0.4 g l^−1^ thiacloprid insecticide granules (Exemptor; Everris). Seedlings were germinated in controlled environment chambers (Reftech, Netherlands) under an 8 h photoperiod at 70% humidity, 20 °C, and photon flux density of 100 µmol m^−2^ s^−1^ of overhead lighting supplied by cool white fluorescent tubes (Reftech, Netherlands). For experiments investigating long-term WUE, seeds were sown within a custom Falcon tube system then stratified. The genotypes that were screened for WUE alterations are identified in Table S2, and all have been described previously. For all experiments, at least two completely independent experimental repeats were performed per genotype and per treatment, with multiple replicate plants within each of the experimental repeats.

### Generation of transgenic lines

To create the *GC1::CCA1:nos* (*GC*)*, GC1::TOC1:nos* (*GT*)*, MYB60::CCA1:nos* (*MC*) and *MYB60::TOC1:nos* (*MT*) constructs, the CaMV *nos* terminator sequence was ligated between the SpeI and NotI restriction sites in the pGREENII0229 binary vector (Hellens et al., 2000). The *GC1* upstream sequence (-1894 to -190) or *MYB60* upstream sequence (-1724 to -429) was then ligated between the KpnI and ApaI restriction sites of pGREENII0229. Finally, the *CCA1* coding sequence or *TOC1* coding sequence, obtained using RT-PCR, was ligated between the restriction XhoI and XmaI sites. Primers used are identified in Table S3. Constructs were transformed into Col-0 wild type Arabidopsis using transformation with *Agrobacterium tumefaciens* strain GV3101. Transformants were identified by screening for phosphinothricin resistance, and then further validated using genomic DNA PCR. Homozygous lines were identified via phosphinothricin (BASTA) resistance, and two independently transformed homozygous lines were investigated in detail per genotype.

Guard cell specificity of promoter activity was investigated using *GC1::GFP:nos* and *MYB60::GFP:nos* promoter-reporter lines (Fig. S3A-C), which were created as above with the *GFP* coding sequence ligated between the XhoI and XmaI restriction sites. Leaf discs (5 mm diameter) from seedlings or mature plants were mounted on microscope slides with dH_2_O, and examined for GFP fluorescence using confocal microscopy (Leica DMI6000). The following settings were used: argon laser at 20% capacity, 488 nm laser at 48% capacity with a bandwidth of 505 nm–515 nm, gain of 1250, offset at 0.2%, 20x or 40x objective, zoom x1 to x4.

### Measurement of water use efficiency

The WUE assay was adapted from Wituszynska et al. (2013) (Wituszyńska et al., 2013). Plants were grown for 6 weeks in modified 50 ml Falcon tubes, under an 8 h photoperiod at 70% humidity, 20 °C, and photon flux density of 100 µmol m^−2^ s^−1^ of overhead lighting supplied by cool white fluorescent tubes (Reftech, Netherlands). The Falcon tube systems consisted of a 50 ml Falcon tube filled with 37.5 ml of a 1:1 ratio of compost: perlite and 35 ml of Milli-Q water (Merck), with the remaining volume filled with a 1:1 ratio of compost: Milli-Q water (Fig. S4). Each Falcon tube lid had a 2 mm diameter hole drilled in its centre to allow plant growth. The lid was spray-painted black (Hycote) because we found that the orange colour of the Falcon tube lid caused leaf curling (Fig. S4). The system was wrapped in aluminium foil to exclude light (Fig. S4). 10-15 seeds were sown through the Falcon tube lid using a pipette. Following stratification, Falcon tube systems were placed under growth conditions using a randomised experimental design. 7 days after germination, seedlings were thinned to one per Falcon tube system, and initial Falcon tube weight was recorded. The seedling-thinning step was sensitive to seedling damage for genotypes with substantially altered morphologies (e.g. *tps1* mutants), reducing the number of replicates available for some genotypes. After 6 weeks of growth, rosette leaf surface area was measured by photography (D50; Nikon) and Fiji software, rosette dry weight was measured (4 d at 60°C), and final Falcon tube weight was recorded. All experiments were stopped before flowering occurred, with the 8 h photoperiod being used to delay flowering as much as possible. Plants were not obviously stressed during the experiment (e.g. leaves did not become purple due to strong anthocyanin accumulation, and plants did not wilt or become contaminated with mildew) (Fig. S4). Negative controls (Falcon tube systems without plants) were used to assess soil water evaporation over 18 experimental batches, with an overall mean weight loss of 0.513 g ± 0.004 g over 6 weeks for plant-free Falcon tubes.

Plant WUE was calculated as follows:

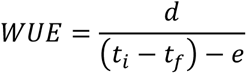

Where *d* is the rosette dry weight at the end of the experiment (mg), *t_i_* and *t_f_* are the falcon tube weight at the start and end of the experiment, respectively (g), and *e* is the amount of water evaporation directly from the compost (g). WUE is derived as mg biomass per ml^−1^ water lost. These calculations assumed that 1 g of weight change was equivalent to a change of 1 ml of water. For examination of individual batches of data (Fig. 1), the WUE of each circadian oscillator genotype was normalized to its respective background to control for variation in the WUE of each background accession and expressed as a percentage of that background. Statistical comparisons with the background lines occurred before normalization. For quantitative investigation of the entire dataset, linear mixed effects models were used (below).

### Measurement of stomatal density

Plants were grown for 7-8 weeks on compost mix. Dental paste (Coltene) was applied to the abaxial surface of fully expanded leaves. Transparent nail varnish (Rimmel) was applied to these leaf moulds once they had set, and then peeled away from the mould using clear adhesive tape (Scotch Crystal). Stomatal and pavement cells were counted within an 800 µm x 800 µm square at the centre of each leaf half, using an epifluorescence microscope (HAL100; Zeiss) and Volocity (Perkin Elmer) and Fiji software. For each experimental repeat, two leaves were sampled per plant and eight plants sampled per genotype. Stomatal index was calculated as follows:

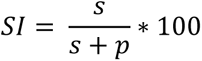

Where *SI* is the stomatal index, *s* the number of stomata in the field of view (800 µm x 800 µm), and *p* the number of pavement cells in the field of view.

### RNA extraction and qRT-PCR

RNA extractions, cDNA synthesis, and qRT-PCR were performed according to (Simon et al., 2018), except approximately 10 seedlings were used per RNA sample and analysis was performed using an MXPro 3005 real time PCR system (Agilent) with 5x HOT FIREPol EvaGreen qPCR mastermix (Solis Biodyne). qRT-PCR primers are provided in Table S4. Rhythmic features within qPCR data were identified using the BioDare2 platform (Zielinski et al., 2014), using the Fast Fourier Transform Non-Linear Least Squares method (FFT-NLLS). One independently-transformed line of each guard cell-specific circadian clock gene overexpressor was also investigated using qRT-PCR conducted on RNA isolated from epidermal peels. Abaxial leaf epidermis was detached, then washed in 10 mM MES (pH 6.15, adjusted using 10 M KOH) to remove RNA derived from ruptured epidermal cells. Each RNA sample was derived from 20 epidermal peels (five plants, four leaves per plant) that were collated and flash-frozen in liquid nitrogen. Guard cell RNA was extracted using the RNeasy UCP Micro Kit (Qiagen) according to manufacturer’s instructions, with the following modification: guard cell lysis was performed by adding glass beads (425 μm - 600 μm diameter, acid washed, from Sigma-Aldrich) and 350 μl RULT buffer to the sample, then vortexed for 5 min.

### Data analysis

Experiments were conducted in a series of 18 separate experimental batches, each including a set of mutants and their corresponding backgrounds. This subdivision ensured high quality experimental attention to each replicate within a large-scale study, and the management of plant growth space. To investigate the nature of any differences between the mutants and their backgrounds in a manner that accounted for between experimental batch-variation in the backgrounds, we used a linear mixed effects modelling approach. Because there are differences between the WUE of Arabidopsis background accessions (Nienhuis et al., 1994; Woodward et al., 2002; Dodd et al., 2004; Masle et al., 2005; Karaba et al., 2007; Ruggiero et al., 2017; Ferguson et al., 2018) (Fig. S2), separate models were generated for each background to avoid comparing accessions with unequal underlying WUE. WUE data were analysed by fitting a linear mixed models using the lme4 package (R package version of lme4 v1.1-21) (Bates et al., 2015) within the R statistical computing platform v3.6.2 (R Core Team, 2019). Using the lmer function, “Mutant” was assigned as a fixed effect (xf) and the “Batch” as a random effect (xr) in order to test for the effect of the mutations on the physiological parameters, whilst controlling for differences between batches. Separate models for each background accession were created with R code:

~~~
model1 <- lmer(Physiological_parameter ~ Mutant + (1|Batch)
~~~

For the dataset from each Arabidopsis background, diagnostic residual plots suggested that the model fits were appropriate for larger datasets (Col-0, C24, L. *er*. and Ws), and for analytical consistency the same model was applied across the entire dataset. Genotypes within two experimental batches (including *toc1*-101 and TOC1-ox) were analysed within a separate model because the backgrounds had rather greater water loss and biomass accumulation than the other experimental batches, so were incorporated into a separate model to obtain the best possible fit. Conditional R^2^ was obtained using the r.squaredGLMM function in the MuMln package v1.43.15 (https://CRAN.R-project.org/package=MuMIn) (Nakagawa and Schielzeth, 2013). The emmeans function (previously lsmeans; (Searle et al., 1980); https://CRAN.R-project.org/package=emmeans); v1.4.3.01) was used to subsequently obtain an estimated marginal mean for each mutant, and conduct post-hoc pairwise comparisons between mutant and its corresponding background, using the Kenward-Roger method for determining degrees of freedom and Tukey method for P value adjustment:

~~~
emmeans(model1, list(pairwise ~ Mutant), adjust = “tukey”)
~~~

With this analysis, this output from group comparisons indicate the statistical significance of any differences of the modelled mutants from the modelled means. This is indicated where relevant on the figures.

### Accession numbers

Arabidopsis Genome Initiative identifiers for the genes mentioned in this study are: *CCA1* (*CIRCADIAN CLOCK ASSOCIATED1*, At2g46830), *CHE* (*CCA1 HIKING EXPEDITION*, At5g08330), *ELF3* (*EARLY FLOWERING3*, At2g25930), *ELF4* (*EARLY FLOWERING4*, At2g40080), *FKF1* (*F BOX1*, At1g68050), *GI* (*GIGANTEA*, At1g22770), *GRP7* (*GLYCINE RICH PROTEIN7*, At2g21660), *KIN10* (*SNF1-RELATED PROTEIN KINASE1.1*, At3g01090), *LKP2* (*LOV KELCH PROTEIN2*, At2g18915), *LUX* (*LUX ARRHYTHMO*, At3g46640), *MYB60* (*MYB DOMAIN PROTEIN60*, At1g08810), *PRR3* (*PSEUDO-RESPONSE REGULATOR3*, At5g60100), *PRR5* (*PSEUDO-RESPONSE REGULATOR5*, At5g24470), *PRR9* (*PSEUDO-RESPONSE REGULATOR9*, At2g46790), *RVE4* (*REVEILLE4*, At5g02840), *TEJ* (*POLY(ADP-RIBOSE)GLYCOHYDROLASE1*, At2g31870), *TIC* (*TIME FOR COFFEE*, At3gt22380), *TOC1* (*TIMING OF CAB EXPRESSION1*, At5g61380), *TPS1* (*TREHALOSE-6-PHOSPHATE SYNTHASE1*, At1g78580), *WNK1* (*WITH NO LYSINE KINASE1*, At3g04910), *ZTL* (*ZEITLUPE*, At5g57360).

## Acknowledgements

We thank Kester Cragg-Barber, James Chen, Ioanna Kostaki, Jean-Charles Isner, Deirdre McLachlan, Peng Sun, Ashutosh Sharma and Dora Cano-Ramirez for technical advice during experimentation. We thank Marc Knight, Tracy Lawson and Steven Penfield for discussions and advice concerning data interpretation and analysis. We thank Keara Franklin, Alex Webb, Paloma Mas, Steve Kay, Isabelle Carre, Takato Imaizumi, Filip Rolland, Ian Graham, Stacey Harmer, and Steven Penfield for donating seed lines. This research was funded the UK Biotechnology and Biological Sciences Research Council (SWBIO DTP awards BB/J014400/1 and BB/M009122/1 and Institute Strategic Programme GEN BB/P013511/1) and the Wolfson Foundation.

