## Supplemental Figures and Tables for "The circadian clock contributes to the long-term water use efficiency of Arabidopsis"

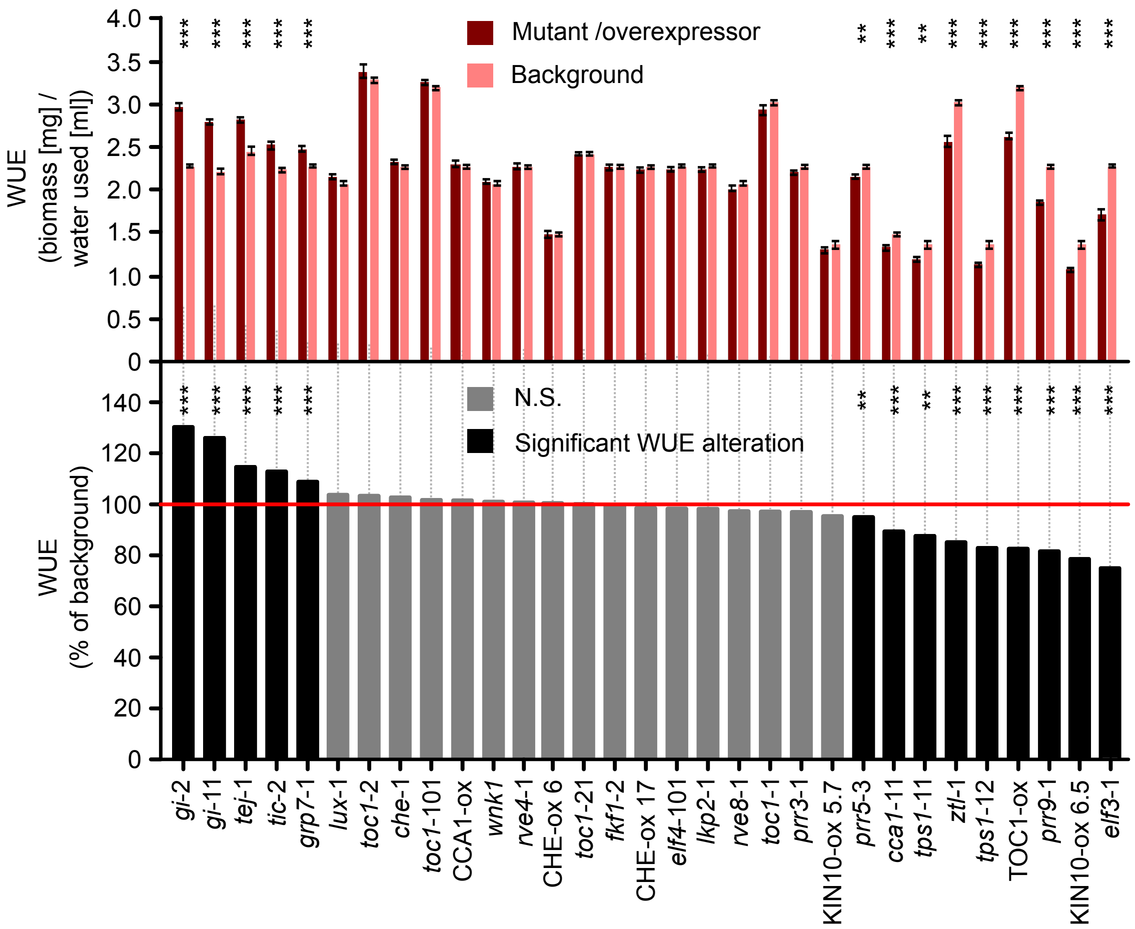


**Figure S1.** The circadian clock regulates long-term water use efficiency of Arabidopsis under light/dark cycles. These data are from a single experimental batch, to summarize the data type used for statistical modelling. The WUE of circadian clock mutants and overexpressors is expressed as absolute WUE and a percentage of their respective background (normalized to the relevant background as 100%, indicated by red reference line). Percentage calculation allows comparison between all genotypes by correcting for WUE variation between background accessions. Data were analysed using independent-samples t-tests, and statistical significance is indicated relative to the background (* = $p < 0.05$; ** = $p < 0.01$; *** = $p < 0.001$; $n = 5 – 15$, ± S.E.M within experimental batch). Statistical analysis was performed on raw WUE data, with significance levels from this indicated on percentage graph also for purposes of comparison.

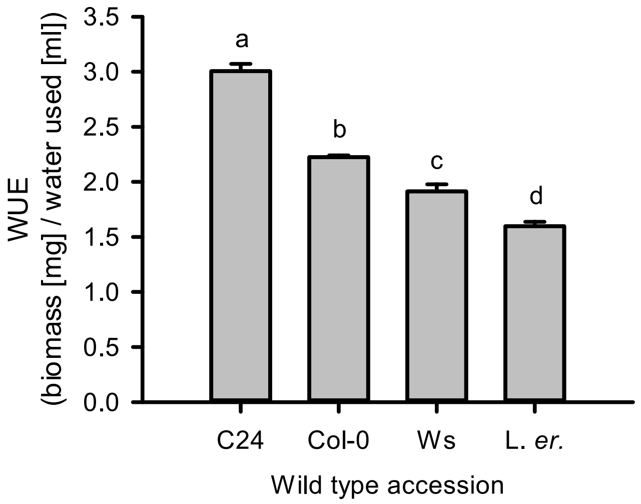


**Figure S2.** Differences in the water use efficiency of wild type accessions. Data were compiled from all independent experimental repeats and analysed with ANOVA and post-hoc Tukey tests. Different letters indicate statistically significant difference ($p < 0.001$; $n$ = 28 - 173; mean ± S.E.M.).

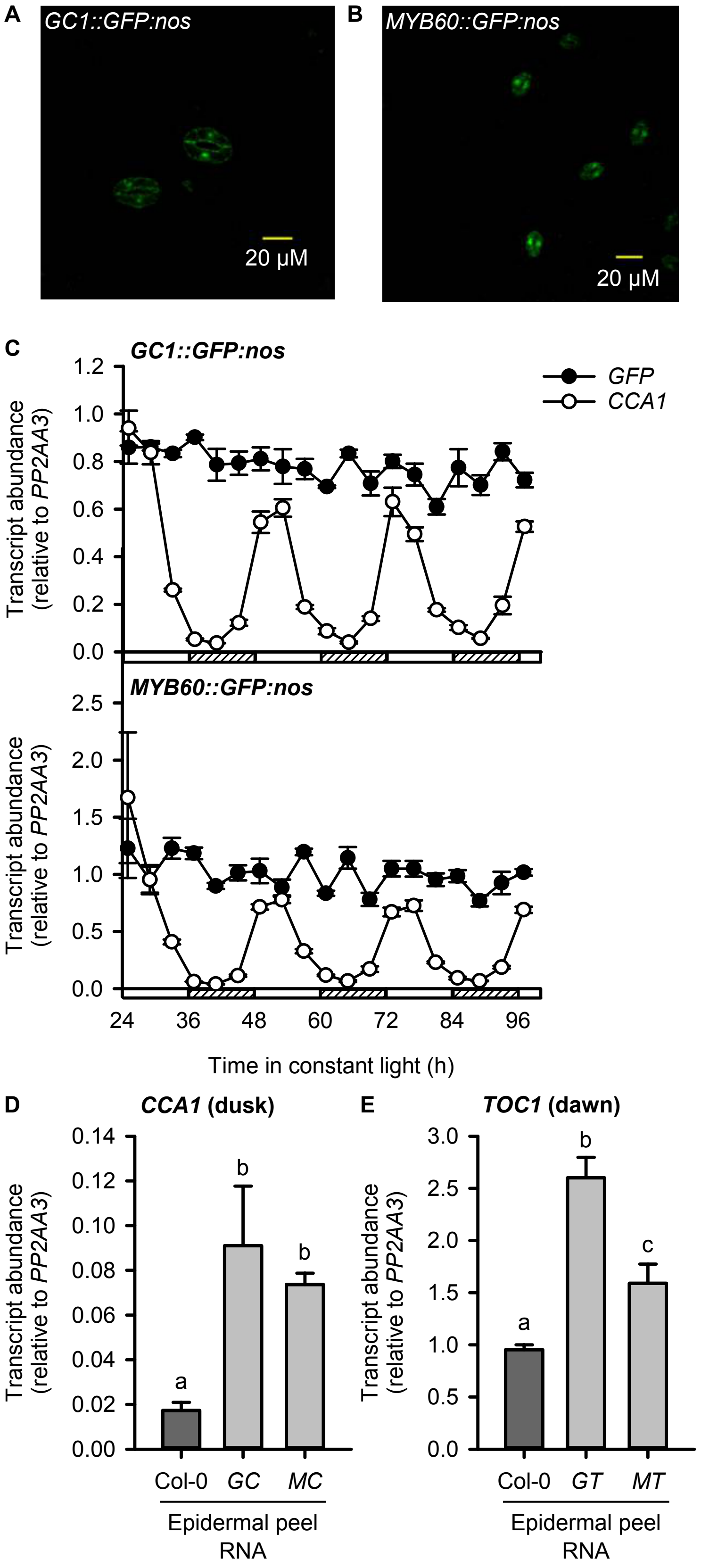


**Figure S3.** Genotyping of plants with guard cell specific overexpression of CCA1 or TOC1. Both (A) *GC1* and (B) *MYB60* promoters have guard cell-specific activity, identified using promoter-*GFP* reporters. Photographs were taken using a confocal microscope with the focal plane on guard cells rather than pavement cells and adjusted identically for brightness and contrast using Fiji. (C) Both *GC1* and *MYB60* promoters have constant, arrhythmic activity, using GFP transcript abundance as a reporter of promoter activity ($n$ = 3, mean ± S.E.M.). *CCA1* transcript abundance was measured concurrently to provide a rhythmic control for comparison. Boxes with and without hashed lines indicate subjective night and day, respectively. *CCA1* transcript abundance was rhythmic for both *GC1::GFP:nos* (FFT-NLLS: period of 24.0 h ± 0.02 h, RAE of 0.47 ± 0.03) and *MYB60::GFP:nos* (FFT-NLLS: period of 24.2 h ± 0.15 h, RAE of 0.58 ± 0.16) seedlings. For both genotypes, FFT-NLLS was unable to fit a waveform to *GFP* transcript abundance. (D, E) Epidermal peel RNA was probed for (D) Relative *CCA1* transcript abundance at dusk for guard cell *CCA1* overexpressors and (E) Relative *TOC1* transcript abundance at dawn for guard cell *TOC1* overexpressors ($n$ = 3; mean ± S.E.M.). We selected these sampling times because they correspond to the approximate lowest times of wild type (D) *CCA1* and (E) *TOC1* transcript abundance in the wild type. Data were analysed with one-way ANOVA ($p < 0.01$) followed by post-hoc Tukey analysis, where different letters indicate statistically significant difference ($p < 0.05$). (C-E) *PP2AA3* was used as the reference transcript. In bar charts, colours indicate the whole plant overexpressor control (black), wild type control (dark grey), and guard cell-specific overexpressor genotypes (light grey).

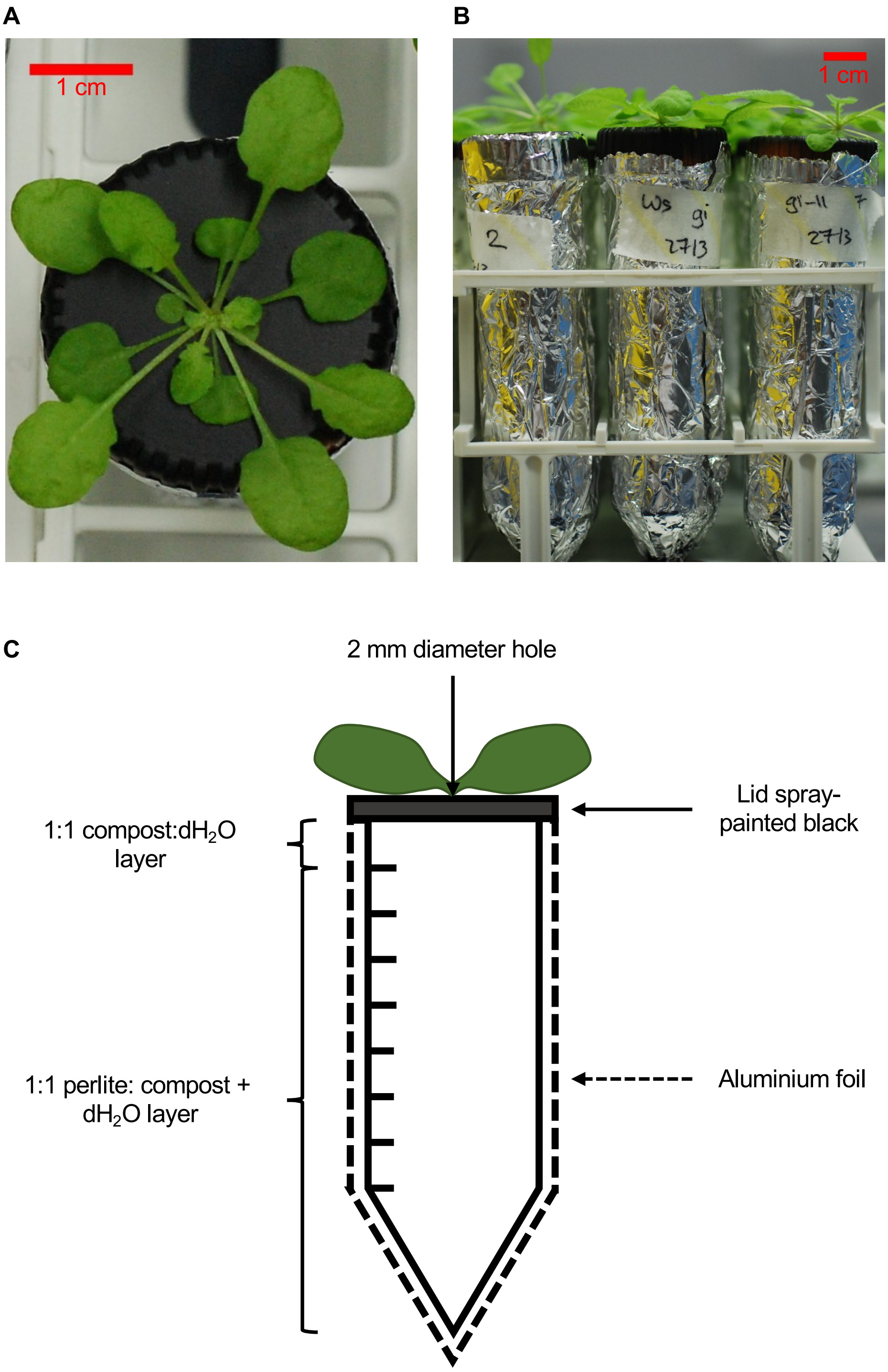


**Figure S4.** Assay used to measure long-term whole plant water use efficiency in Arabidopsis. (A, B) Representative photographs of the optimised WUE assay setup from (A) above and (B) the side. (C) Each falcon tube contained a bottom layer of 1:1 (volume) perlite:compost with dH$_2$O, and an upper layer of 1:1 compost:dH$_2$O (volume). (A-C) Falcon tube lids were spray-painted black and have a 2 mm diameter centrally-located hole through which the plant grows. The entire falcon tube system was wrapped in aluminium foil. Further details are provided in Materials and Methods.

**Table S1:** Quality of fit of linear mixed effects models to experimental data. Separate models were formed for each background line because the WUE of each background differed considerably (Fig. S2). Water use and biomass gain of the backgrounds for two experimental batches differed substantially from the other experimental batches, so separate models were run for these batches to obtain a better model fit and estimated mean.

| **Physiological parameter** | **Background line within model** | **Conditional $R^2$** |
|---|---|---|
| WUE | Col-0 | 0.908 |
| | L. *er*. | 0.917 |
| | Ws | 0.914 |
| | C24 | 0.737 |
| | C24 (*tej*-1 background) | 0.687 |
| Water used | Col-0 | 0.718 |
| | Col-0 (experimental batches 23 and 24) | 0.414 |
| | L. *er*. | 0.813 |
| | Ws | 0.801 |
| | C24 | 0.684 |
| | C24 (*tej*-1 background) | 0.265 |
| Dry biomass gain | Col-0 | 0.708 |
| | Col-0 (experimental batches 23 and 24) | 0.551 |
| | L. *er*. | 0.813 |
| | Ws | 0.815 |
| | C24 | 0.687 |
| | C24 (*tej*-1 background) | 0.201 |

**Table S2: Arabidopsis genotypes screened for water use efficiency.**

| Gene ID (AGI code) | Gene | Genotype | Background | Reference |
|---|---|---|---|---|
| AT2G46830 | *CCA1* | *cca1*-11 | Ws | Hall et al. (2003) |
| AT2G46830 | *CCA1* | CCA1-ox | Col | Wang and Tobin (1998) |
| AT5G08330 | *CHE* | *che*-1<br>*che*-2<br>*CCA1::LUC*+ | Col | Pruneda-Paz et al. (2009) |
| AT5G08330 | *CHE* | CHE-ox<br>*CCA1::LUC*+ 17 | Col | Pruneda-Paz et al. (2009) |
| AT5G08330 | *CHE* | CHE-ox<br>*CCA1::LUC*+ 6 | Col | Pruneda-Paz et al. (2009) |
| AT2G25930 | *ELF3* | *elf3*-1 | Col | Zagotta et al. (1992) |
| AT2G40080 | *ELF4* | *elf4*-101 | Col | Khanna et al. (2003) |
| AT1G68050 | *FKF1* | *fkf1*-2 | Col | Imaizumi et al. (2003) |
| AT1G22770 | *GI* | *gi*-11<br>*CCA1::LUC* | Ws | Ding et al. (2007) |
| AT1G22770 | *GI* | *gi*-2 | Col | Fowler et al. (1999) |
| AT2G21660 | *CCR2* | *grp7*-1 | Col | Streitner et al. (2008) |

| | | | | |
|---|---|---|---|---|
| AT3G01090 | *KIN10* | KIN10-ox 5.7 | L. *er*. | Baena-Gonzalez et al. (2007) |
| AT3G01090 | *KIN10* | KIN10-ox 6.5 | L. *er*. | Baena-Gonzalez et al. (2007) |
| AT2G18915 | *LKP2* | *lkp2*-1 | Col | Imaizumi et al. (2005) |
| AT3G46640 | *LUX* | *lux*-1 | Col | Hazen et al. (2005) |
| AT5G60100 | *PRR3* | *prr3*-1 | Col | Michael et al. (2003) |
| AT5G24470 | *PRR5* | *prr5*-3 *TOC1::LUC* | Col | Michael et al. (2003) |
| AT2G46790 | *PRR9* | *prr9*-1 | Col | Eriksson et al. (2003) |
| AT5G02840 | *RVE4* | *rve4*-1 | Col | Alonso et al. (2003) |
| AT3G09600 | *RVE8* | *rve8*-1 | Col | Rawat et al. (2011) |
| AT2G31870 | *TEJ* | *tej*-1 | C24 | Panda et al. (2002) |
| AT3G22380 | *TIC* | *tic*-2 | Col | Ding et al. (2007) |
| AT5G61380 | *TOC1* | *toc1*-1 *CAB2::LUC* | C24 | Millar et al. (1995) |

| | | | | |
|---|---|---|---|---|
| AT5G61380 | *TOC1* | *toc1*-2 *CAB2::LUC* | C24 | Strayer et al. (2000) |
| AT5G61380 | *TOC1* | *toc1*-21 *CAB2::LUC* | Ws | Ding et al. (2007) |
| AT5G61380 | *TOC1* | *toc1*-101 | Col | Kikis et al. (2005) |
| AT5G61380 | *TOC1* | *TOC1*-ox | Col | Más et al. (2003) |
| AT1G78580 | *TPS1* | *tps1*-11 | L. *er*. | Gomez et al. (2010) |
| AT1G78580 | *TPS1* | *tps1*-12 | L. *er*. | Gomez et al. (2010) |
| AT1G78580 | *TPS1* | *tps1*-13 | L. *er*. | Gomez et al. (2010) |
| AT3G04910 | *WNK1* | *wnk1* | Col | Alonso et al. (2003) |
| AT5G57360 | *ZTL* | *ztl*-1 *CAB2::LUC* | C24 | Somers et al. (2000) |

**Table S3: Primers used for cloning**

| Gene ID (AGI code) of target | Product | Primer sequences (5’ – 3’) |
|---|---|---|
| AT1G22690 | *GC1* promoter sequence with 5’ KpnI and 3’ ApaI restriction sites | F: TCGACGGTACCGAGTAAAGATTCAGTAACCCGA<br>R: ACTTAGGGCCCGTGATTTTGAAGTAGTGTGTGA |
| AT1G08810 | *MYB60* promoter sequence with 5’ KpnI and 3’ ApaI restriction sites | F: ATTAAGGTACCCACAA GGACACAAGGACATA<br>R: AATATTGGGCCCCTCTTCCTCTAGATCTCTCTGAG |
| AT2G46830 | *CCA1* coding sequence with 5’ XhoI and 3’ XmaI restriction sites | F:ATCGACCTCGAGATGGAGACAAATTCGTCTGG<br>R:GTCAGACCCGGGAAAATAGAGTCTCATGTGGAAGC |
| AT5G61380 | *TOC1* coding sequence with 5’ XhoI and 3’ XmaI restriction sites | F: ATCGACCTCGAGATGGATTTGAACGGTGAGTGTA<br>R: AGGAAGCCCGGGTCAAG TTCCCAAAGCATCATC |

| | | |
|---|---|---|
| | *GFP* coding sequence with 5’ XhoI and 3’ XmaI restriction sites | F:ATCAGACTCGAGATGAGTAAAGGAGAAGAACTTTTC<br>R: ATCGAGCCCGGGTTATT TGTATAGTTCATCCATGC |
| | *nos* terminator sequence with 5’ SpeI and 3’ NotI restriction sites | F: ACTAGTGAATTTCCCCGATCGTTC<br>R: GCGGCCGCTCTAGTAACATAGATG |

**Table S4: Primers used for qRT-PCR**

| Gene ID (AGI code) of target | Transcript | Primer sequences (5' – 3') |
|---|---|---|
| AT2G46830 | *CCA1* | F: GCACTTTCCGCGAGTTCTTG<br>R: TGACTCCTTTCTTACCCTGTTATTCTG |
| | *GFP* | F: CCATCTTCTTCAAGGACGAC<br>R: CCTTAAGCTCGATCCTGTTG |
| AT1G13320 | *PP2AA3* | F: TAACGTGGCCAAAATGATGC<br>R: GTTCTCCACAACCGCTTGGT |
| AT5G61380 | *TOC1* | F: TCTTCGCAGAATCCCTGTGAT<br>R: GCTGCACCTAGCTTCAAGCA |
